# Diversity and abundance of ring nucleases in type III CRISPR-Cas loci

**DOI:** 10.1101/2024.09.24.614671

**Authors:** Ville Hoikkala, Haotian Chi, Sabine Grüschow, Shirley Graham, Malcolm F. White

**Affiliations:** School of Biology, University of St Andrews, St Andrews, KY16 9ST, UK

## Abstract

Most type III CRISPR-Cas systems facilitate immune responses against invading mobile genetic elements such as phages by generating cyclic oligoadenylates (cOAs). Downstream effectors activated by cOAs are typically non-specific proteins that induce damage to essential cellular components, thereby preventing phage epidemics. Due to these toxic effects, it is crucial that the production and concentration of cOAs remain under tight regulatory control during infection-free periods or when deactivating the immune response after clearing an infection. Type III CRISPR loci often encode enzymes known as ring nucleases (RNs) that bind and degrade specific cOAs, while some effectors are auto-deactivating. Despite the discovery of several classes of RNs, a comprehensive bioinformatic analysis of type III CRISPR-Cas loci in this context is lacking. Here, we examined 38,742 prokaryotic genomes to provide a global overview of type III CRISPR loci, focussing on the known and predicted RNs. The candidate RNs Csx16 and Csx20 are confirmed as active enzymes, joining Crn1-3. Distributions and patterns of co-occurrence of RNs and associated effectors are explored, allowing the conclusion that a sizeable majority of type III CRISPR systems regulate cOA levels by degrading the signalling molecules, which has implications for cell fate following viral infection.

## Introduction

CRISPR-Cas is an adaptive prokaryotic immune system that incorporates genomic fragments of invading mobile genetic elements (MGEs) into the host’s chromosomal CRISPR array (1,2). These fragments, called spacers, are expressed during subsequent infections as CRISPR RNA (crRNA) and help interference complexes to target invading nucleic acids via sequence complementarity (3). Type III CRISPR-Cas systems use multi-subunit interference complexes, hallmarked by Cas10 – a large protein that can harbour two main functions (3). While a minority of Cas10 proteins use an HD nuclease domain to cleave ssDNA non-specifically (4,5), over 90% of Cas10s have a cyclase domain that generates second messenger signalling molecules (6–9). The signal molecules produced by Cas10, either cyclic oligoadenylates (cOA) or SAM-AMP (10), accumulate in the cell and activate downstream effector proteins that are typically encoded by genes in the same operon. The effector proteins contain a sensory domain that captures the signal, leading to the allosteric activation of the effector domain that may be a nuclease, protease, or a membrane-disrupting domain (see reviews (11,12)). Activation of effectors thus tends to have toxic consequences for cells and their actions can lead to growth arrest. This outcome is sometimes called abortive infection; an umbrella term under which diverse immune mechanisms are often grouped, all sharing the ecological rationale of sacrificing the host for the sake of the population (see review (13)). Whether phage defence mechanisms actually lead to cell death *in vivo* is debated (14), and the presence of ring nucleases (RNs) in CRISPR type III systems provides a mechanism to avoid this outcome.

RNs are small proteins that degrade cOA signal molecules, thus thwarting the activation of downstream effectors. They are generally encoded in CRISPR-Cas loci and are also utilized by viruses as anti-CRISPR (Acr) proteins (15). It is not currently known whether their primary function in cells is to deactivate defence in an ongoing infection state or to curtail signal molecule levels in a non-infected state. To date, three families of RNs have been experimentally verified. The first RN to be discovered, Crn1 (CRISPR-associated ring nuclease 1), was identified biochemically from a lysate of *Saccharolobus solfataricus* (16). Crn1 is a metal-independent phosphodiesterase that binds cA_4_ in its dimeric CARF domains and degrades it into linear A_2_>P (cyclic 2’,3’ phosphate) products. Crn1 is most commonly found in crenarchaeal type III CRISPR loci and has a relatively low cA_4_ cleavage rate, which may be tuned to remove cA_4_ from the cell without disrupting the immune response (16).

Crn2 and its virally-encoded counterpart AcrIII-1 are also cA_4_-cleaving RNs (15). This class of RN is characterised by the DUF1874 domain, which forms a dimeric cA_4_ recognition domain. AcrIII-1 degrades cA_4_ much more quickly than Crn1, consistent with its role in subverting type III CRISPR signalling during viral infection. This family of RNs has also been found fused to the bacterial effector Csx1, constituting a self-limiting cA_4_-activated ribonuclease (17).

Crn3, previously known as Csx3, constitutes the third family of cA_4_-cleaving RNs. The structure of Crn3 is only distantly related to the CARF superfamily, harbouring closer resemblance to sulfate transporter and anti-sigma factor antagonist (STAS) domains (18–20). Crn3 binds cA_4_ by sandwiching the molecule between two adjacent dimers that tetramerise in a head-to-tail orientation (18). The phosphodiesterase reaction is manganese dependent and generates linear A_2_-P products. Crn3 is sometimes fused to a AAA-ATPase domain of unknown function (20,21).

In addition to the dedicated RNs, several type III CRISPR effectors have been shown to degrade their cOA activators within the binding site of the sensory domain, effectively functioning as self-limiting enzymes. All cA_6_-dependent Csm6 family ribonucleases studied to date display cA_6_ RN activity in the CARF recognition domain (22–24). Likewise cA_4_-activated Csm6/Csx1 effectors are also capable of degrading their activator within the CARF domain (25–27). Recently, the CalpL effector, which uses a SAVED domain to bind cA_4_, has also been confirmed as a RN (28,29).

The CRISPR-associated Csx15, Csx16 and Csx20 proteins have been predicted to be RNs, based on sequence and genomic neighbourhood analyses (20), but this has not yet been experimentally confirmed. RNs are thus common in type III CRISPR systems but have not been studied systematically. Here, we undertook an extensive analysis of type III CRISPR loci, mapping the known and predicted RNs and exploring their association with the diverse range of effector proteins. The cA_4_ RN activities of Csx15, Csx16 and Csx20 are investigated biochemically and structural modelling is used to predict their mechanisms of cA_4_ recognition.

## Methods

### Data preparation

A total of 38,742 complete bacterial and archaeal genomes were downloaded from NCBI on March 25^th^ 2024. For phage genomes, the May 2024 set of 28,114 curated genomes from the Millard database (30) was downloaded.

### Bioinformatic methods

A previously described Snakemake (31) pipeline was used as the basis for type III CRISPR locus characterisation (6). In short, the pipeline uses custom-built Cas10 HMM profiles to find type III CRISPR loci, which are further analysed using CCTyper (32) and a panel of HMM profiles to discover effector proteins. The pipeline was modified to enable ring nuclease detection and phage genome analysis.

Custom-built HMM-libraries were constructed using published sequence data in NCBI for ring nucleases Crn1, Crn2 and Crn3. For Csx15, Csx16 and Csx20, libraries were built based on previously published sequences (20), followed by expansion of homologs through blastp searches against the NCBI protein database. Each protein coding sequence with 6 kbp of a type III CRISPR locus was analysed for RNs using hmmscan from the Hmmer 3.3.2 package (33). To prevent cross-annotation with effectors that have similar domains (e.g. CARF), a maximum length cutoff of 250 aa for RNs during annotation.

The steps for constructing the Cas10 phylogenetic tree were outlined in (6). In short, each Cas10 was annotated for the presence of cyclase or nuclease domains. The Cas10 sequences were then aligned with Muscle (34). A phylogenetic tree file was built using FastTree2 (35) with arguments -wag and -gamma, and visualised in RStudio 2024.4.0.735 (36) using ggtree (37). The tree was annotated with CRISPR subtype data from CCTyper (in a few cases corrected manually) and with the presence of known and candidate ring nuclease families. The signal molecule associated with each locus was inferred from the type(s) of effector present and annotated for each locus in the tree.

A network interaction graph was made with Gephi (https://gephi.org/) using RN/effector co-occurrence data. Co-occurrences between effectors and between RNs were removed to highlight those between RNs and effectors.

To search for ring nucleases in phage genomes, the proteomes of all 28,114 phage genomes in the Millard phage database were analysed using our RN HMM libraries using Hmmer 3.3.2 (33) similar to the type III CRISPR-Cas analysis outlined above.

### Cloning, expression and purification of Csx15, 16 and 20

Synthetic genes (Supplementary Table 1) encoding Csx15, Csx16 and Csx20, codon optimised for expression in *E. coli*, were purchased from Integrated DNA Technologies (IDT), Coralville, USA, cloned into the pEHisV5TEV vector (38) between the *Nco*I and *Bam*HI sites and transformed into DH5α cells. Construct integrity was confirmed by sequencing (Eurofins Genomics, DE).

The constructs were transformed into *E. coli* C43 (DE3) and proteins were expressed according to the standard protocols previously described (38). In brief, 2 litres of culture were induced with 0.4 mM isopropyl-β-D-1-thiogalactoside (IPTG) at an OD_600_ of ∼0.8 and grown for 4 h or overnight at 25 °C. Cells were harvested (4000 rpm; Beckman Coulter JLA-8.1 rotor) and resuspended in lysis buffer and lysed by sonication. Proteins were purified with an immobilised metal affinity chromatography (IMAC) column (HisTrapFF, Cytiva, Marlborough, USA), washed with 5 column volumes (CV) of loading buffer and eluted with a linear gradient of loading buffer plus 0.5 M imidazole. Following his-tag removal by TEV protease, proteins were subjected to a second IMAC step and the unbound fraction collected. Size exclusion chromatography was used to further purify the proteins, which were eluted isocratically as described previously (38). Pure proteins were concentrated, aliquoted and stored frozen at -70 °C.

### Ring nuclease activity of Csx15, Csx16 and Csx20

Ring nuclease activity was assayed by incubating 1 µM of each protein with 100 µM synthetic cA_3_, cA_4_ or cA_6_ (Biolog) in reaction buffer (20 mM Tris-HCl pH7.5, 250 mM NaCl and 5 mM EDTA) at 30 °C for 60 min. The reaction was quenched by adding methanol and vortexing. The mixture was then dried, before resuspension in water for HPLC analysis on an UltiMate3000 UHPLC system (Thermo Fisher scientific) with a C18 column (Kinetex EVO 2.1 × 50 mm, particle size 2.6 µm). The column temperature was set at 40 °C and absorbance was monitored at 260 nm. Samples were analysed by gradient elution with solvent A (20 mM ammonium acetate, pH 8.5) and solvent B (methanol) as a flow rate of 0.3 ml/min as follows: 0-0.5 min, 1% B; 0.5-6 min, 1-15% B; 6-7 min, 100% B.

## Results

### Structural models and RN activity of Csx15, Csx16 and Csx20

We showed previously that 92% of type III CRISPR loci likely function via nucleotide signalling, with active Cas10 polymerase domains (6). Of these, the predominant signalling molecule is cA_4_, so it is perhaps unsurprising that the three stand-alone RNs identified to date, Crn1-3, are all specific for cA_4_ (15,16,18). Csx15, 16 and 20 were previously identified as candidate RNs based on genome context, sequence conservation and structural predictions (20). However, these predictions have not been confirmed biochemically, so we first sought to test this specific hypothesis.

The structures of Csx15, Csx16 and Csx20 were modelled as dimers using Alphafold3 (AF3) (39) (Figure 1A). Two AMP ligands were included to help predict cOA binding sites. Inclusion of these ligands improved the ordering of the mobile loops that probably become structured on cOA binding. All three models were predicted with high confidence (ptm / iptm/ranking scores 0.91/0.90/0.94, 0.79/0.75/0.77 and 0.95/0.94/0.95 for Csx15, 16 and 20, respectively). To gain more insight into the likely cOA binding sites of the proteins, we mapped conserved residues from a diverse range of homologues for each protein (Supplementary Figure 1) onto the AF3 structures (Supplementary Figure 2). This revealed a cluster of conserved residues on one face of Csx16 and Csx20 that correspond with the modelled AMP binding sites and likely pinpoint the binding site for cOA. However, for Csx15, conserved residues were more prevalent on the “bottom” face of the dimer, away from the canonical cOA binding site. This situation is reminiscent of Crn3, which forms filaments of head-to-tail dimers sandwiching cA_4_ (18). Structural comparisons using the DALI server (40) yielded hits for Csx15 with SAVED and CARF family proteins (Z-score >5 for PDB accessions 7RWM, 8Q3Z, 8FMF and 7QDA); Csx16 yielded a significant hit (Z-score 4.8) with PDB 2J6B – a viral homologue of Crn2 (41) whilst Csx20 yielded a strong match (Z-score 8.0) with the Crn1 family protein Sso2081 (PDB 7YGH) (42). Pairwise comparison of the predicted structures of Csx16 and Csx20 yielded a DALI Z-score of 8.9, consistent with a common core fold consisting of a 5-stranded β-sheet flanked by α-helices. These structural features are consistent with a derived Rossman fold (CARF-like) structure for Csx15, Csx16 and Csx20 (20).

**Figure 1.**
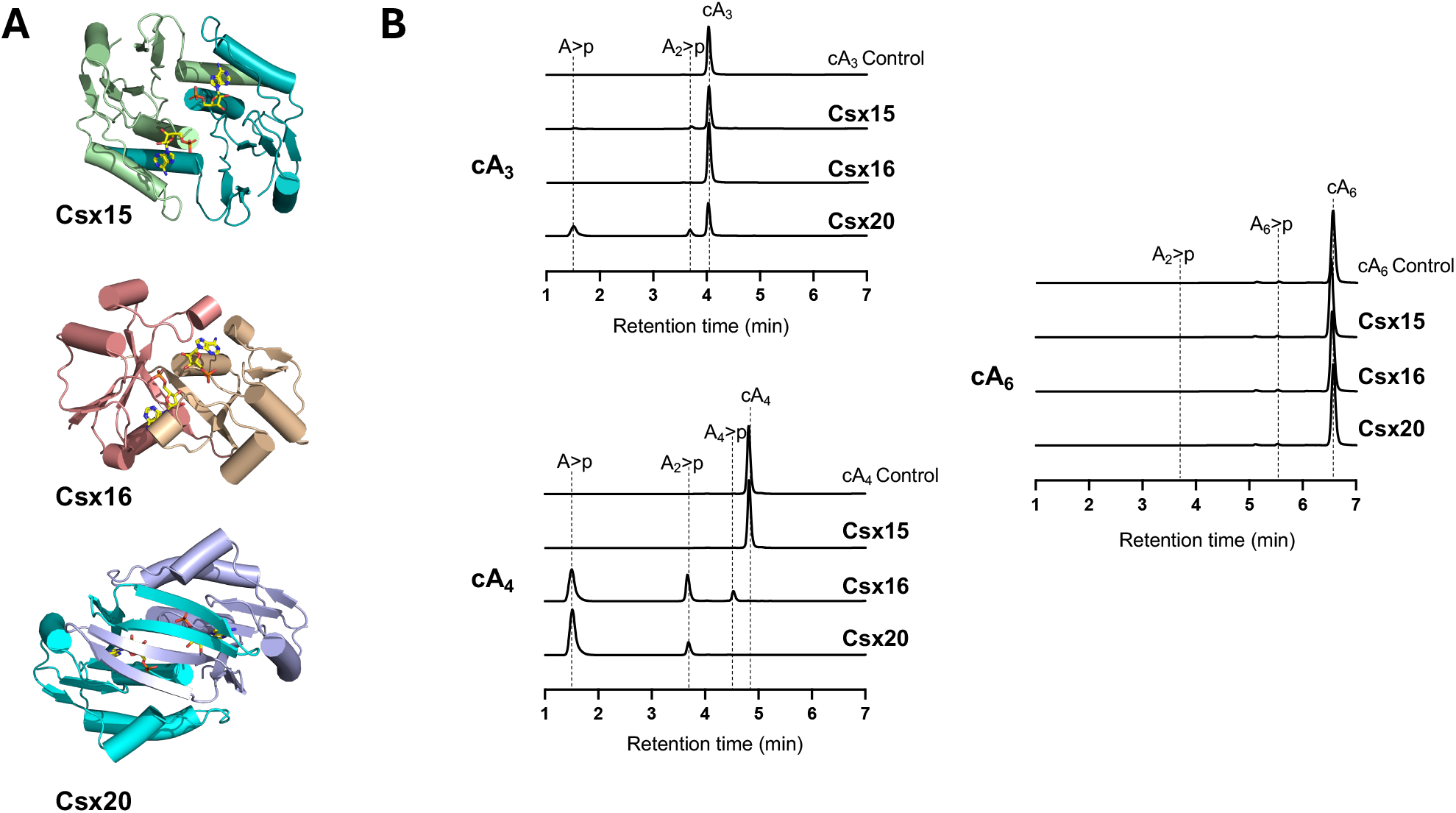
Structural modelling and RN activity of Csx15, Csx16 and Csx20. **A**. Dimeric protein AF3 models are shown with 2 AMP molecules (yellow sticks) modelled to mimic the cOA binding site, subunits are coloured differently for ease of interpretation. **B**. RN activity of Csx15, Csx16 and Csx20 against cA_3_, cA_4_ and cA_6_, monitored by HPLC. Csx16 and Csx20 degrade cA_4_ into linear products. Standards and characterised reaction products are labelled. (>p represents a 2’,3’-cyclic phosphate).

Representative examples of Csx15, Csx16 and Csx20 were cloned, expressed in *E. coli* and purified to homogeneity as described in the methods (Supplementary Figure 3A). The proteins were tested individually for the ability to degrade cA_3_, cA_4_ and cA_6_ under conditions of 100-fold substrate excess for 1 h (Figure 1B). Csx16 and Csx20 showed clear RN activity against cA_4_, fully degrading it to small linear products. Neither enzyme degraded cA_6_, whilst only very minor activity for Csx20 against cA_3_ could be observed. Csx15, on the other hand, displayed no RN activity against any cOA species (Figure 1B). To explore RN activity more fully, we repeated these experiments with a ten-fold higher concentration of protein (10 µM) (Supplementary Figure 3B). Under these conditions, Csx20 fully degraded cA_3_ as well as cA_4_, whilst Csx16 retained specificity for cA_4_. Very limited activity of Csx15 against cA_3_ and cA_6_, but not cA_4_, could be observed. We conclude that Csx16 and Csx20 are cA_4_-specific RNs, whilst the function of Csx15 remains uncertain. It is possible that we have not found the correct reaction conditions to reveal the activity of Csx15. Alternatively, it could conceivably act as a cOA “sponge” rather than a RN, as phage-encoded sponge proteins are known to sequester cyclic nucleotides and inhibit cellular defences (43,44).

### Distribution and co-occurrence patterns of ring nucleases

Having explored the RN activity of Csx15, 16 and 20, we proceeded to analyse the distribution of RNs across the Cas10 tree (Figure 2). This analysis immediately demonstrated that RNs are widespread in Cas10 loci, and when found in an effector-containing locus (96% of RN instances), they are associated only with cA_4_-dependent effectors. In the 698 CRISPR loci that contained a cA_4_-activated effector, 40% had an associated RN. The most frequently observed RN was Csx20 (80 loci), followed by Csx16 (69 loci), Crn1 (44 loci), Crn3 (38 loci), Csx15 (18 loci) and Crn2 (7 loci). There was no obvious bias in the distribution of RNs with respect to CRISPR subtypes that utilise cOA signalling, while loci from subtypes III-C and III-F, which lack an active Cas10 cyclase, have very few associated RNs. Csm6 and Csm6-2, which are activated by cA_6_, are not found in association with known RNs, which may reflect intrinsic RN activity by these effectors.

**Figure 2.**
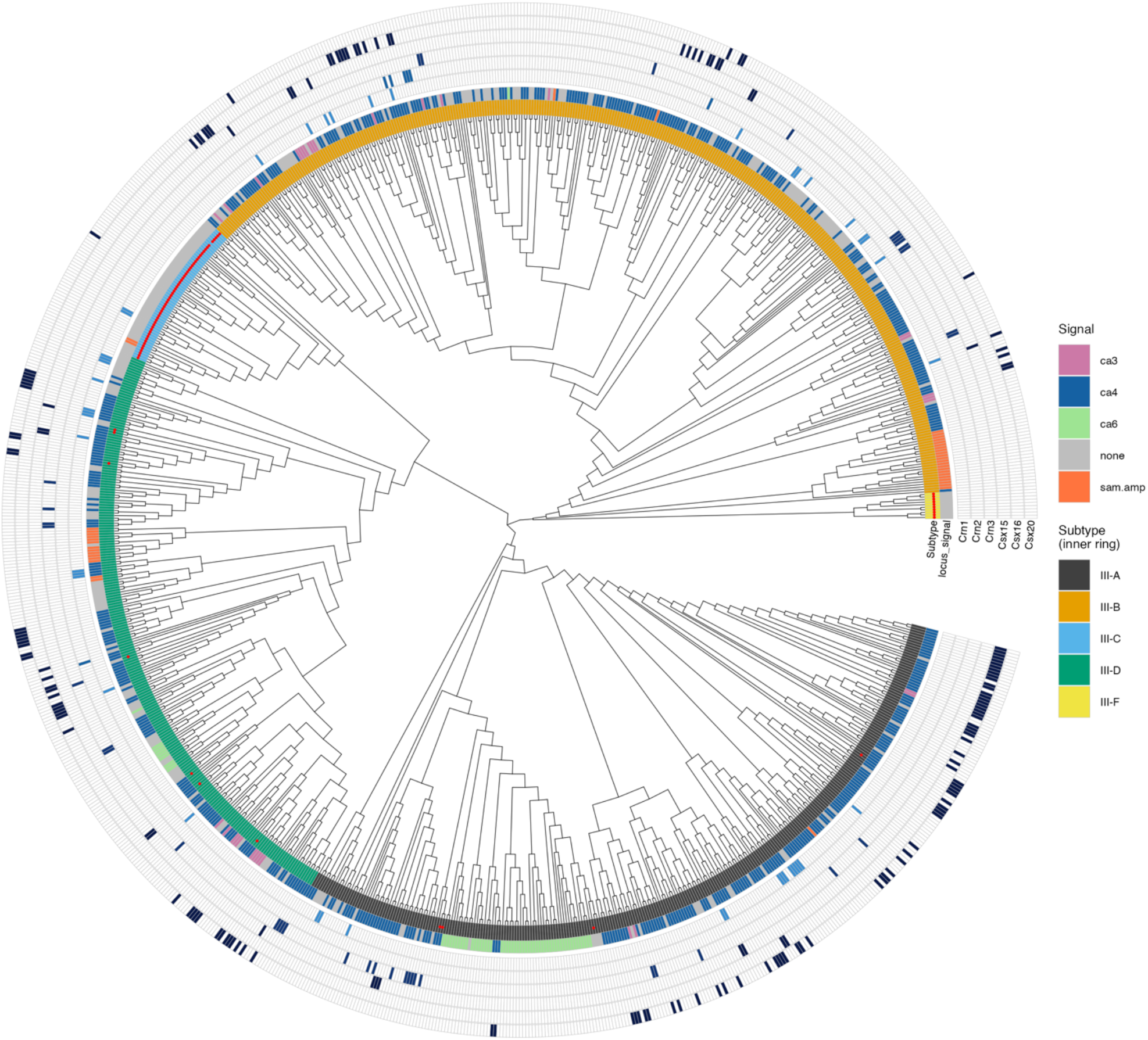
Phylogenetic tree of Cas10 with ring nuclease distribution. The signalling molecule used by each system is predicted based on effector content of the locus, as defined in (6), and coloured according to the key. Instances of candidate ring nucleases Crn1, Crn2, Crn3, Csx15, Csx16 and Csx20 are indicated in concentric circles. Cas10s lacking a clear active cyclase domain are indicated by a red dot in the subtype ring.

Only 10 of the 246 RN-positive loci (∼4%) contained more than one RN, which supports the hypothesis that they are all performing equivalent roles. An exception to this rule was Csx15, which co-occurs with Crn1 in 7 loci and as the sole RN in 11 loci. We investigated the frequency of Csx15 across all ∼40 000 RefSeq genomes and found that Csx15 is found in genomes without a type III CRISPR-Cas locus in 25% of cases. Along with the peculiar co-occurrence pattern with Crn1, this suggests Csx15 may not be a canonical RN.

In 32 cases, RNs were found in loci with no effector, suggesting either the presence of unknown effectors in the locus or a role for RNs *in trans* by another type III CRISPR locus with a known effector. In support of the latter possibility, 9 of the 32 effector-lacking loci are co-located in a genome with a separate type III locus that contains an effector – this count is 0 for the 214 loci that include both an effector and a RN.

The co-occurrence of ring nucleases and effectors is shown in Figure 3A. All characterised ring nucleases preferentially degrade cA_4_, so it is unsurprising (but reassuring) that they co-occur with cA_4_-specific effectors. The most abundant effector in our dataset, Csx1, defined here as the cA_4_-activated family of CARF-HEPN dimeric effector proteins (6) is found in association with each of the ring nucleases analysed here and has a particularly strong association with Csx20. We also analysed loci with a single known effector to quantify the proportion that encoded a RN (Figure 3B). The observation that only 34% of Csx1-containing loci encode a known RN may be at least partially explained by the fact that some Csx1 family members have intrinsic RN activity (25,27). In support of this, the Can1-2 effector family (45–47), which lack intrinsic RN activity, are associated with RNs in 66% of loci whereas for Cami1, which possesses RN activity (48,49), the proportion is only 30%. Following this logic, the observation that the membrane bound effector Cam1 is found alongside RNs in 81% of loci suggests that this effector also lacks intrinsic RN activity, although that has yet to be tested (50). None of the characterised cA_4_-specific RNs are found associated with effectors that use a different signal molecule. While some of these effectors, such as Csm6, are effective RNs in their own right (22–24,27), others, such as NucC, are not (51). This could suggest that RNs targeting molecules other than cA_4_ simply await discovery, or alternatively that infection outcomes are different in these cases.

**Figure 3.**
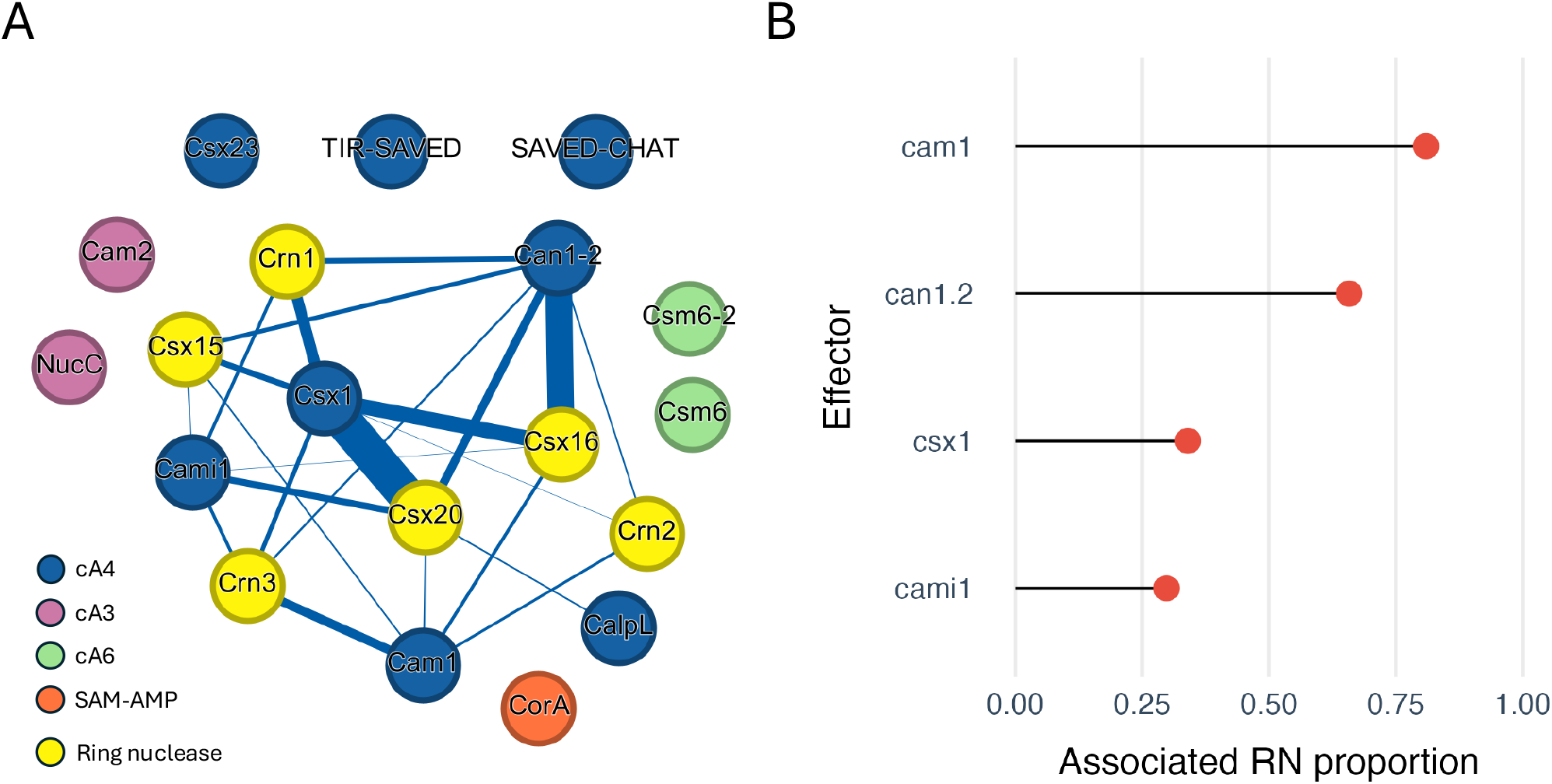
Co-occurrence and fusion of ring nucleases and effectors in type III CRISPR loci. **A**. Gephi plot showing co-occurrence patterns between effectors and ring nucleases. Ring nucleases are in yellow, cA_3_, cA_4_ and cA_6_-activated effectors in orange, pink and blue, respectively and SAM-AMP-activated effectors in green. **B**. The proportion of effector instances that are associated with a RN (data only includes loci with one effector and zero or one RNs).

Phage genomes were also investigated for ring nucleases. Among the 28,114 phage genomes in the Millard database, the previously described anti-CRISPR protein AcrIII-1 (the phage-encoded homolog of Crn2) (15) was by far the most abundant ring nuclease, present in 36 genomes. Additionally, Csx20 was found in five genomes and Csx16 in two genomes. The high relative abundance of AcrIII-1/Crn2 in phage genomes compared to its presence in only 7 prokaryotic genomes supports the view that it likely originated from viruses as an anti-CRISPR and was subsequently adopted by cellular hosts to regulate cOA levels.

We released an interactive web portal for visualizing and filtering the bioinformatic results. The website is available at https://vihoikka.github.io/rn_browser.

## Discussion

By confirming the RN activity of Csx16 and 20, we have expanded the catalogue of extrinsic RNs to five families. These are widely distributed across the type III CRISPR systems that signal via cA_4_ – the predominant signalling molecule used by 84% of all known effector instances (6). When combined with the observation that intrinsic RN activity is frequently observed in CARF or SAVED sensor domains of effectors, we can conclude with some confidence that an “off-switch” is an important component of most type III CRISPR systems. Exceptions, such as the prophage-encoded systems described in *Vibrio* species that use NucC or Csx23 effectors (51,52), exist but likely represent a small minority.

The prokaryotic immune system has two major components that potentiate an antiviral response by generation of cOA species: type III CRISPR-Cas and CBASS (Cyclic oligonucleotide-based antiphage signalling system) (53). Each system responds to infection by activating a specialised cyclase that generates a cyclic nucleotide second messenger which in turn activates one or more effector proteins to provide an immune response. CBASS and type III CRISPR-Cas have much in common, with shared signalling molecules such as cA_3_ and effectors such as NucC (51,54) and TIR-SAVED (6,55). Each uses the signal amplification possible with a second messenger to activate a toxic response to infection that slows down viral replication at the cost of cell fitness. Their main point of divergence appears to be the fact that type III CRISPR systems often encode an “off-switch” while, in contrast, CBASS activation appears to be a “one way street” that may often result in cell death or growth arrest (56).

This fundamental difference between type III CRISPR and CBASS defence, despite all their similarities, may reflect the fact that the former is activated early in infection (by viral mRNA) whilst the latter is viewed as a “last ditch” defence, typically activated late in the infection cycle (for example, by intracellular phage capsid proteins (43)) when other defences have failed (57,58). CBASS defence, functioning at the level of herd immunity, is clearly worth having, as the system is fairly common in bacteria (59). Nevertheless, cells appear to avoid altruistic suicide when possible. One significant caveat is that RNs may primarily function to prevent aberrant activation of type III CRISPR-Cas defence (ie activation of Cas10 by non-viral RNA), rather than to clear up after a *bona fide* infection. The growth-arrested state facilitated by type III CRISPR-Cas may also enable other defence systems to clear the infecting virus, which could then be followed by reversal of the arrested state by RNs. Such cooperation between defence systems has been found between the RNA-targeting type VI CRISPR-Cas systems and restriction modification (RM), where restriction enzymes cleave phage genome whilst the cell is under Cas13-induced dormancy (60). Looking ahead, there is a pressing need for studies that tackle these questions using an evolutionary, population-based approach.

Although we have analysed six candidate RN families that account for a large proportion of the cA_4_-signalling CRISPR systems, it is possible and, indeed, likely that further RN families remain to be discovered. The function of Csx15 also remains an open question at this point. By looking closely at the loci which lack a known RN and which have an effector with intrinsic RN activity, we hope to unveil new RN families in the future.

## Acknowledgements

This work was supported by a European Research Council Advanced Grant (Grant REF 101018608 to MFW). HC acknowledges the support of the China Scholarship Council (code 202008420207).

## Data accessibility statement

The Snakemake pipeline and accompanying scripts are available for review at https://github.com/vihoikka/ring_nucleases. This repository also contains the HMM profiles for Cas10s, effectors and all six ring nuclease families in this study. An interactive website for browsing the bioinformatic results is available at https://vihoikka.github.io/rn_browser.

---

**Supplementary Figure 1.**
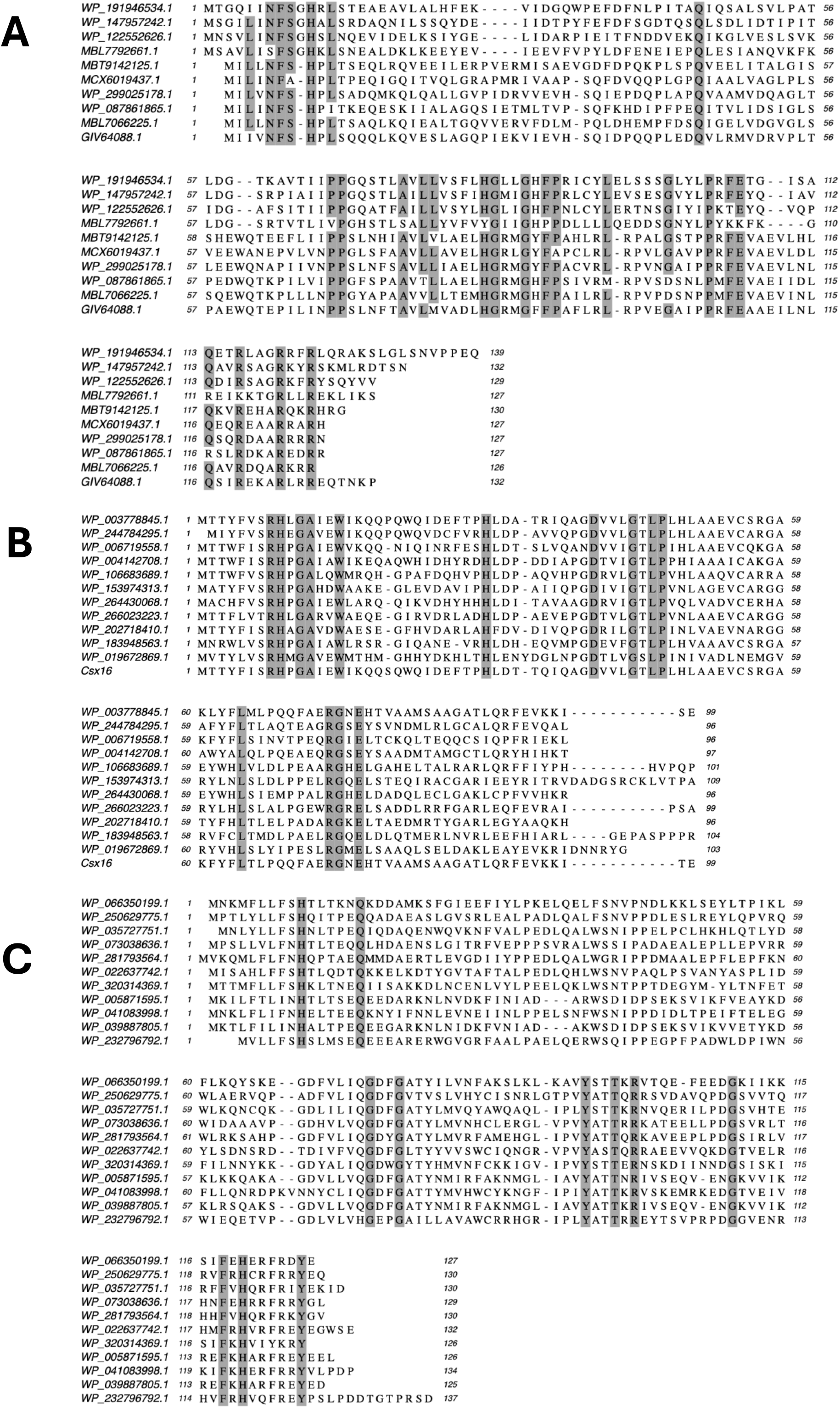
Multiple sequence alignments of selected orthologues of Csx15 (A), Csx16 (B) and Csx20 (C), with conserved residues shaded.

**Supplementary Figure 2.**
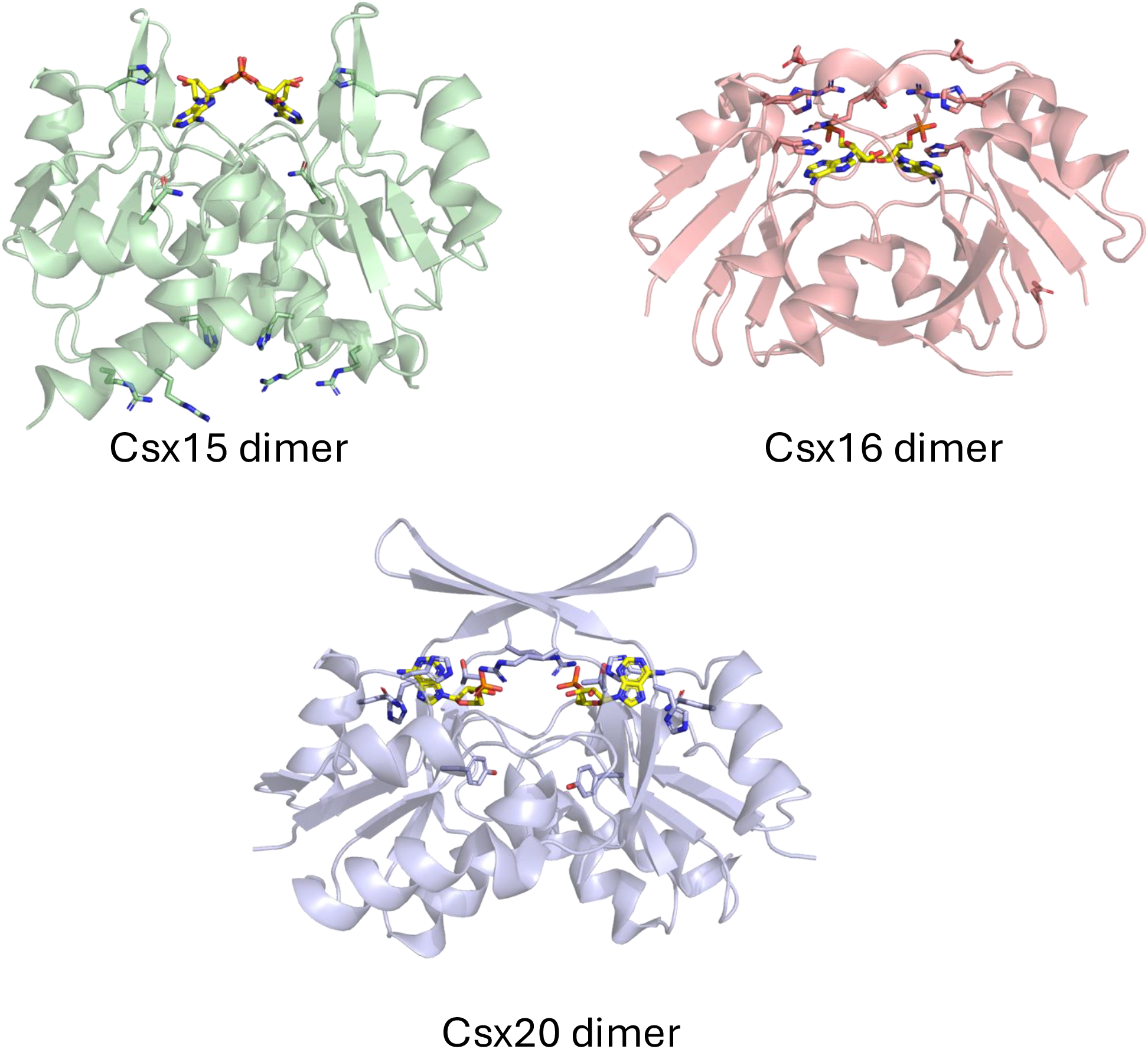
AF3 dimer models of Csx15, 16 and 20, showing the side chains of conserved residues as sticks and 2 AMP moieties in the presumed cA_4_ binding site.

**Supplementary Figure 3.**
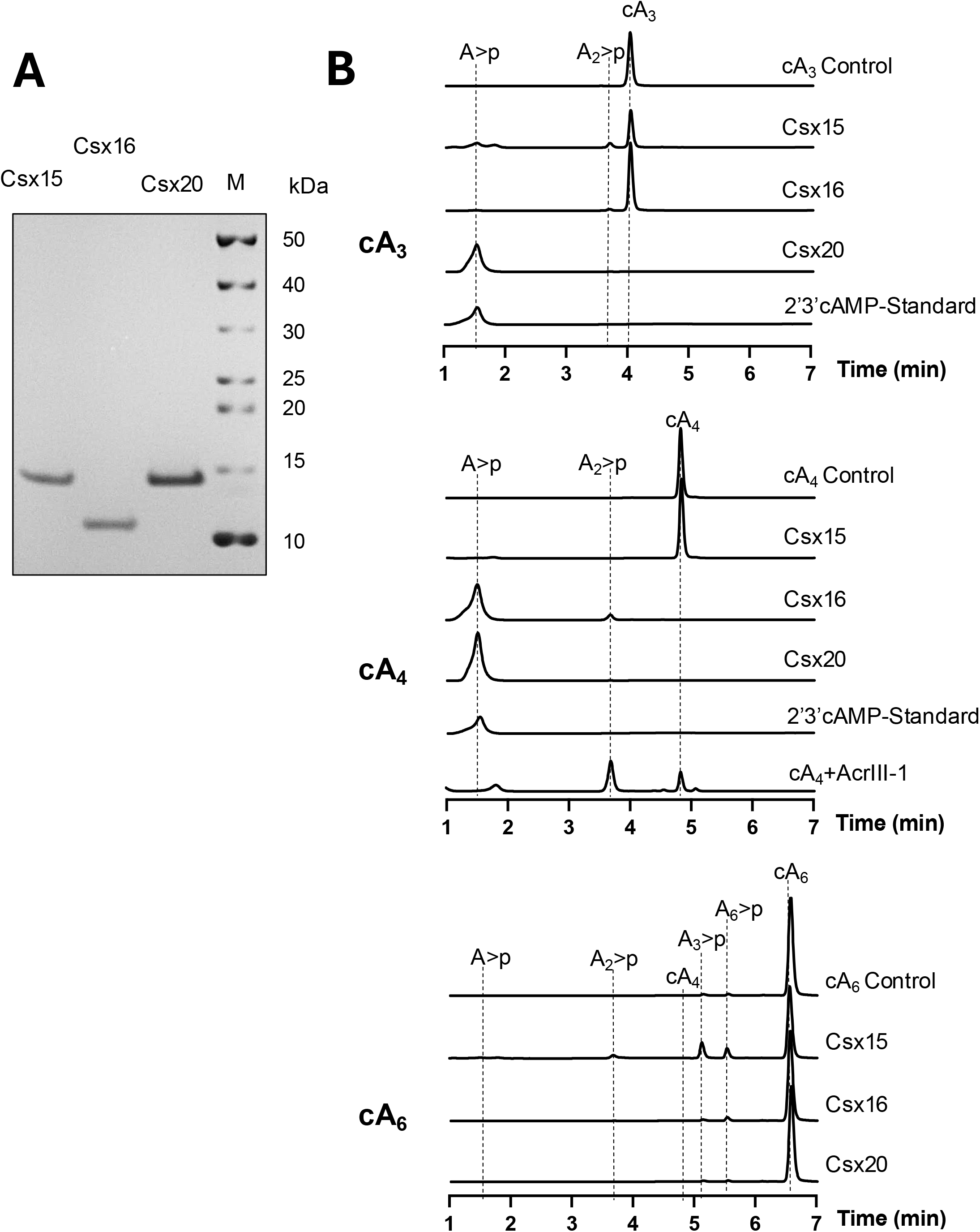
**A**. SDS-PAGE analysis of purified Csx15, Csx16 and Csx20. **B**. HPLC analysis of RN activity of Csx15, Csx16 and Csx20 (10 µM protein; 70 µM cOA species, 1 h). Standards are indicated along with the previously characterised reaction products of AcrIII-1 incubated with cA_4_ (15).

**Supplementary Table 1.**
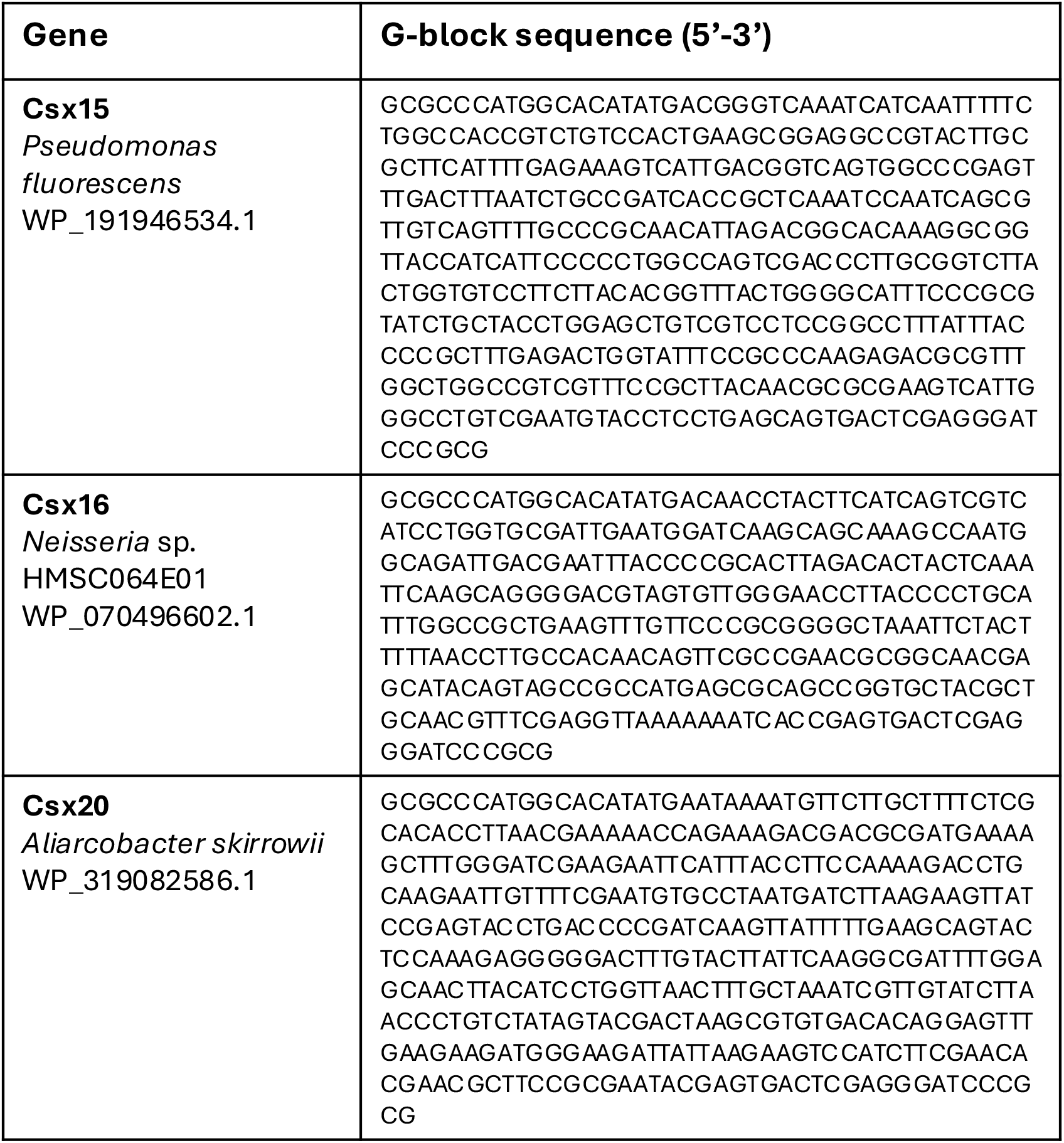
Synthetic gene sequences for Csx15, Csx16 and Csx20

## References

1. Barrangou R, Fremaux C, Deveau H, Richards M, Boyaval P, Moineau S, et al. CRISPR provides acquired resistance against viruses in prokaryotes. Science. 2007 Mar;315(5819):1709–12.

2. Mojica FJM, Díez-Villaseñor C, García-Martínez J, Soria E. Intervening Sequences of Regularly Spaced Prokaryotic Repeats Derive from Foreign Genetic Elements. J Mol Evol. 2005;60(2):174–82.

3. Makarova KS, Wolf YI, Alkhnbashi OS, Costa F, Shah SA, Saunders SJ, et al. An updated evolutionary classification of CRISPR–Cas systems. Nat Rev Microbiol. 2015 Sep;13(11):722–36.

4. Jung TY, An Y, Park KH, Lee MH, Oh BH, Woo E. Crystal structure of the Csm1 subunit of the Csm complex and its single-stranded DNA-specific nuclease activity. Struct Lond Engl 1993. 2015 Apr 7;23(4):782–90.

5. Samai P, Pyenson N, Jiang W, Goldberg GW, Hatoum-Aslan A, Marraffini LA. Co-transcriptional DNA and RNA cleavage during type III CRISPR-Cas immunity. Cell. 2015 May 21;161(5):1164–74.

6. Hoikkala V, Graham S, White MF. Bioinformatic analysis of type III CRISPR systems reveals key properties and new effector families. Nucleic Acids Res. 2024 May 29;gkae462.

7. Kazlauskiene M, Kostiuk G, Venclovas Č, Tamulaitis G, Siksnys V. A cyclic oligonucleotide signaling pathway in type III CRISPR-Cas systems. Science. 2017;357(6351):605–9.

8. Niewoehner O, Garcia-Doval C, Rostøl JT, Berk C, Schwede F, Bigler L, et al. Type III CRISPR–Cas systems produce cyclic oligoadenylate second messengers. Nature. 2017 Aug;548(7669):543–8.

9. Wiegand T, Wilkinson R, Santiago-Frangos A, Lynes M, Hatzenpichler R, Wiedenheft B. Functional and Phylogenetic Diversity of Cas10 Proteins. CRISPR J. 2023 Apr;6(2):152–62.

10. Chi H, Hoikkala V, Grüschow S, Graham S, Shirran S, White MF. Antiviral type III CRISPR signalling via conjugation of ATP and SAM. Nature. 2023 Oct;622(7984):826–33.

11. Athukoralage JS, White MF. Cyclic Nucleotide Signaling in Phage Defense and Counter-Defense. Annu Rev Virol. 2022 Sep 29;9(1):451–68.

12. Steens JA, Salazar CRP, Staals RHJ. The diverse arsenal of type III CRISPR-Cas-associated CARF and SAVED effectors. Biochem Soc Trans. 2022 Oct 31;50(5):1353–64.

13. Lopatina A, Tal N, Sorek R. Abortive Infection: Bacterial Suicide as an Antiviral Immune Strategy. Annu Rev Virol. 2020;7(1):1–14.

14. Fernández-García L, Wood TK. Phage-Defense Systems Are Unlikely to Cause Cell Suicide. Viruses. 2023 Aug 24;15(9):1795.

15. Athukoralage JS, McMahon SA, Zhang C, Grüschow S, Graham S, Krupovic M, et al. An anti-CRISPR viral ring nuclease subverts type III CRISPR immunity. Nature. 2020;577(7791):572–5.

16. Athukoralage JS, Rouillon C, Graham S, Grüschow S, White MF. Ring nucleases deactivate type III CRISPR ribonucleases by degrading cyclic oligoadenylate. Nature. 2018;562(7726):277–80.

17. Samolygo A, Athukoralage JS, Graham S, White MF. Fuse to defuse: a self-limiting ribonuclease-ring nuclease fusion for type III CRISPR defence. Nucleic Acids Res. 2020;48(11):6149–56.

18. Athukoralage JS, McQuarrie S, Grüschow S, Graham S, Gloster TM, White MF. Tetramerisation of the CRISPR ring nuclease Crn3/Csx3 facilitates cyclic oligoadenylate cleavage. eLife. 2020;9:e57627.

19. Brown S, Gauvin CC, Charbonneau AA, Burman N, Lawrence CM. Csx3 is a cyclic oligonucleotide phosphodiesterase associated with type III CRISPR-Cas that degrades the second messenger cA4. J Biol Chem. 2020 Oct 30;295(44):14963–72.

20. Makarova KS, Timinskas A, Wolf YI, Gussow AB, Siksnys V, Venclovas Č, et al. Evolutionary and functional classification of the CARF domain superfamily, key sensors in prokaryotic antivirus defense. Nucleic Acids Res. 2020;48(16):8828–47.

21. Shah SA, Alkhnbashi OS, Behler J, Han W, She Q, Hess WR, et al. Comprehensive search for accessory proteins encoded with archaeal and bacterial type III CRISPR-cas gene cassettes reveals 39 new cas gene families. RNA Biol. 2019 Apr 3;16(4):530–42.

22. Garcia-Doval C, Schwede F, Berk C, Rostøl JT, Niewoehner O, Tejero O, et al. Activation and self-inactivation mechanisms of the cyclic oligoadenylate-dependent CRISPR ribonuclease Csm6. Nat Commun. 2020 Mar 27;11(1):1596.

23. McQuarrie S, Athukoralage JS, McMahon SA, Graham S, Ackermann K, Bode BE, et al. Activation of Csm6 ribonuclease by cyclic nucleotide binding: in an emergency, twist to open. Nucleic Acids Res. 2023 Oct 27;51(19):10590–605.

24. Smalakyte D, Kazlauskiene M, F Havelund J, Rukšėnaitė A, Rimaite A, Tamulaitiene G, et al. Type III-A CRISPR-associated protein Csm6 degrades cyclic hexa-adenylate activator using both CARF and HEPN domains. Nucleic Acids Res. 2020;48(16):gkaa634.

25. Athukoralage JS, Graham S, Grüschow S, Rouillon C, White MF. A Type III CRISPR Ancillary Ribonuclease Degrades Its Cyclic Oligoadenylate Activator. J Mol Biol. 2019 Jul 12;431(15):2894–9.

26. Du L, Zhu Q, Lin Z. Molecular mechanism of allosteric activation of the CRISPR ribonuclease Csm6 by cyclic tetra-adenylate. EMBO J. 2024 Jan;43(2):304–15.

27. Jia N, Jones R, Yang G, Ouerfelli O, Patel DJ. CRISPR-Cas III-A Csm6 CARF Domain Is a Ring Nuclease Triggering Stepwise cA4 Cleavage with ApA>p Formation Terminating RNase Activity. Mol Cell. 2019 Sep 5;75(5):944–956.e6.

28. Binder SC, Schneberger N, Schmitz M, Engeser M, Geyer M, Rouillon C, et al. The SAVED domain of the type III CRISPR protease CalpL is a ring nuclease. Nucleic Acids Res. 2024 Aug 21;gkae676.

29. Smalakyte D, Ruksenaite A, Sasnauskas G, Tamulaitiene G, Tamulaitis G. Filament formation activates protease and ring nuclease activities of CRISPR SAVED-Lon. bioRxiv; 2024 [cited 2024 Aug 13]. p. 2024.05.08.593097. Available from: https://www.biorxiv.org/content/10.1101/2024.05.08.593097v1

30. Cook R, Brown N, Redgwell T, Rihtman B, Barnes M, Clokie M, et al. INfrastructure for a PHAge REference Database: Identification of Large-Scale Biases in the Current Collection of Cultured Phage Genomes. PHAGE. 2021 Dec;2(4):214–23.

31. Mölder F, Jablonski KP, Letcher B, Hall MB, Tomkins-Tinch CH, Sochat V, et al. Sustainable data analysis with Snakemake [Internet]. F1000Research; 2021 [cited 2024 Aug 12]. Available from: https://f1000research.com/articles/10-33

32. Russel J, Pinilla-Redondo R, Mayo-Muñoz D, Shah SA, Sørensen SJ. CRISPRCasTyper: Automated Identification, Annotation, and Classification of CRISPR-Cas Loci. CRISPR J. 2020 Dec 1;3(6):462–9.

33. Eddy SR. Accelerated Profile HMM Searches. PLoS Comput Biol. 2011 Oct;7(10):e1002195.

34. Edgar RC. Muscle5: High-accuracy alignment ensembles enable unbiased assessments of sequence homology and phylogeny. Nat Commun. 2022 Nov 15;13(1):6968.

35. Price MN, Dehal PS, Arkin AP. FastTree 2 – Approximately Maximum-Likelihood Trees for Large Alignments. PLOS ONE. 2010 Mar 10;5(3):e9490.

36. RStudio Team. RStudio: Integrated Development for R. RStudio [Internet]. Boston, MA: RStudio, PBC; 2020. Available from: http://www.rstudio.com/

37. Yu G. Using ggtree to Visualize Data on Tree-Like Structures. Curr Protoc Bioinforma. 2020 Mar;69(1):e96.

38. Rouillon C, Athukoralage JS, Graham S, Grüschow S, White MF. Investigation of the cyclic oligoadenylate signaling pathway of type III CRISPR systems. Methods Enzymol. 2019;616:191–218.

39. Abramson J, Adler J, Dunger J, Evans R, Green T, Pritzel A, et al. Accurate structure prediction of biomolecular interactions with AlphaFold 3. Nature. 2024 Jun;630(8016):493–500.

40. Holm L. Dali server: structural unification of protein families. Nucleic Acids Res. 2022 Jul 5;50(W1):W210–5.

41. Keller J, Leulliot N, Cambillau C, Campanacci V, Porciero S, Prangishvili D, et al. Crystal structure of AFV3-109, a highly conserved protein from crenarchaeal viruses. Virol J. 2007 Jan 22;4(1):12.

42. Du L, Zhang D, Luo Z, Lin Z. Molecular basis of stepwise cyclic tetra-adenylate cleavage by the type III CRISPR ring nuclease Crn1/Sso2081. Nucleic Acids Res. 2023 Mar 21;51(5):2485–95.

43. Huiting E, Cao X, Ren J, Athukoralage JS, Luo Z, Silas S, et al. Bacteriophages inhibit and evade cGAS-like immune function in bacteria. Cell. 2023 Feb 16;186(4):864–876.e21.

44. Jenson JM, Li T, Du F, Ea CK, Chen ZJ. Ubiquitin-like conjugation by bacterial cGAS enhances anti-phage defence. Nature. 2023 Apr;616(7956):326–31.

45. McMahon SA, Zhu W, Graham S, Rambo R, White MF, Gloster TM. Structure and mechanism of a Type III CRISPR defence DNA nuclease activated by cyclic oligoadenylate. Nat Commun. 2020 Jan 24;11(1):500.

46. Rostøl JT, Xie W, Kuryavyi V, Maguin P, Kao K, Froom R, et al. The Card1 nuclease provides defence during type III CRISPR immunity. Nature. 2021;590(7847):624–9.

47. Zhu W, McQuarrie S, Grüschow S, McMahon SA, Graham S, Gloster TM, et al. The CRISPR ancillary effector Can2 is a dual-specificity nuclease potentiating type III CRISPR defence. Nucleic Acids Res. 2021;49(5):2777–89.

48. Mogila I, Tamulaitiene G, Keda K, Timinskas A, Ruksenaite A, Sasnauskas G, et al. Ribosomal stalk-captured CARF-RelE ribonuclease inhibits translation following CRISPR signaling. Science. 2023 Dec;382(6674):1036–41.

49. Rouillon C, Schneberger N, Chi H, Blumenstock K, Da Vela S, Ackermann K, et al. Antiviral signalling by a cyclic nucleotide activated CRISPR protease. Nature. 2023 Feb;614(7946):168–74.

50. Baca CF, Yu Y, Rostøl JT, Majumder P, Patel DJ, Marraffini LA. The CRISPR effector Cam1 mediates membrane depolarization for phage defence. Nature. 2024 Jan 10;1–8.

51. Grüschow S, Adamson CS, White MF. Specificity and sensitivity of an RNA targeting type III CRISPR complex coupled with a NucC endonuclease effector. Nucleic Acids Res. 2021 Dec 16;49(22):13122–34.

52. Grüschow S, McQuarrie S, Ackermann K, McMahon S, Bode BE, Gloster TM, et al. CRISPR antiphage defence mediated by the cyclic nucleotide-binding membrane protein Csx23. Nucleic Acids Res. 2024 Apr 12;52(6):2761–75.

53. Millman A, Melamed S, Amitai G, Sorek R. Diversity and classification of cyclic-oligonucleotide-based anti-phage signalling systems. Nat Microbiol. 2020;5(12):1608–15.

54. Lau RK, Ye Q, Birkholz EA, Berg KR, Patel L, Mathews IT, et al. Structure and Mechanism of a Cyclic Trinucleotide-Activated Bacterial Endonuclease Mediating Bacteriophage Immunity. Mol Cell. 2020;77(4):723–733.e6.

55. Hogrel G, Guild A, Graham S, Rickman H, Grüschow S, Bertrand Q, et al. Cyclic nucleotide-induced helical structure activates a TIR immune effector. Nature. 2022 Aug;608(7924):808–12.

56. Duncan-Lowey B, Kranzusch PJ. CBASS phage defense and evolution of antiviral nucleotide signaling. Curr Opin Immunol. 2022 Feb 1;74:156–63.

57. Cohen D, Melamed S, Millman A, Shulman G, Oppenheimer-Shaanan Y, Kacen A, et al. Cyclic GMP– AMP signalling protects bacteria against viral infection. Nature. 2019;574(7780):691–5.

58. Krüger L, Gaskell-Mew L, Graham S, Shirran S, Hertel R, White MF. Reversible conjugation of a CBASS nucleotide cyclase regulates bacterial immune response to phage infection. Nat Microbiol. 2024 Apr 8;1–14.

59. Tesson F, Hervé A, Mordret E, Touchon M, d’Humières C, Cury J, et al. Systematic and quantitative view of the antiviral arsenal of prokaryotes. Nat Commun. 2022 May 10;13(1):2561.

60. Williams MC, Reker AE, Margolis SR, Liao J, Wiedmann M, Rojas ER, et al. Restriction endonuclease cleavage of phage DNA enables resuscitation from Cas13-induced bacterial dormancy. Nat Microbiol. 2023 Mar;8(3):400–9.

